# Development and characterization of infectious clones of two strains of Usutu virus

**DOI:** 10.1101/2020.08.05.238543

**Authors:** Tyler Bates, Christina Chuong, Seth A. Hawks, Pallavi Rai, Nisha K. Duggal, James Weger-Lucarelli

## Abstract

Usutu virus (USUV; genus *Flavivirus*; family *Flaviviridae*) is a mosquito-borne, positive-sense RNA virus that is currently causing significant die-offs in numerous bird species throughout Europe and has caused infections in humans. Currently, there are no molecular clones for USUV, hence, hindering studies on the pathogenesis and transmission of USUV. In this report, we demonstrate the development and characterization of infectious clones for two modern strains of USUV isolated from Europe and Africa. We show that the infectious clone-derived viruses replicated similarly to the parental strains in both mammalian and insect cells. Additionally, we observed similar levels of replication and pathogenesis in two mouse models. This reverse genetics system will aid the scientific community in studying and developing USUV infection, transmission, diagnostics, and vaccines.

## Introduction

Usutu virus (USUV; genus *Flavivirus*; family *Flaviviridae)* is an emerging arthropod-borne virus, or arbovirus, currently causing major fatalities in birds throughout Europe, especially in Eurasian blackbirds — *Turdus merula* (1). It was first isolated in 1959 in South Africa from *Culex neavei* mosquitoes (2), and since then has spread to Europe, circulating between avian and mosquito species (3). The virus has been detected in Italy, Greece, France, Spain, Poland, Hungary, Czech Republic, Serbia, the United Kingdom, Croatia, the Netherlands, Switzerland, and Germany (as reviewed by (4)). In Germany alone, USUV has caused the death of an estimated 420,000 Eurasian blackbirds (5). This virus causes sporadic spillover cases in humans who are generally asymptomatic; however, a subset of those infected people may experience fever, rash, and, in severe cases, meningoencephalitis (6-8).

USUV threatens to become a public health crisis, given its genetic and ecological similarities to West Nile virus (WNV), which likewise spread from Africa to Europe, and then to the USA in 1999 (9). Similar to the other members of the Japanese encephalitis virus (JEV) serocomplex— JEV, WNV, and St. Louis encephalitis virus (SLEV)—USUV circulates among *Culex* mosquitoes and various bird species, while humans and other mammals serve as dead-end hosts (10-12). A majority of USUV infections, like other members of the JEV serocomplex, are asymptomatic; however, some infected people may develop febrile illness, including fever, nausea, headaches, and vomiting. In more severe cases, USUV, like other JEV serocomplex viruses, can cause encephalitis, meningitis, or meningoencephalitis in humans (13-15). Despite the detrimental impact on bird populations and the threat to public health, no molecular tools for studying USUV currently exist. To this end, we developed infectious cDNA clones for two modern USUV strains and characterized their use *in vitro* and *in vivo*.

Infectious clones are useful tools for studying viral pathogenesis, replication, transmission, evolution, and for vaccine development (16). The current lack of infectious clones to study USUV hinders progress towards increasing our understanding of its fundamental biology and identifying critical viral genetic determinants of pathogenesis and transmission. In this report, we describe infectious clones for two recently isolated USUV strains—TM-Netherlands 2016 and UG09615 2010—collected in the Netherlands and Uganda, respectively, and performed phenotypic comparisons to the parental viruses in both *in vitro* and *in vivo* models. We first used bacteria to generate the infectious clones but found they were attenuated due to mutations likely introduced during bacterial propagation. We then used an innovative bacteria-free approach to correct these mutations and demonstrate that these clone-derived viruses behave phenotypically identical to the parental strains *in vitro* and *in vivo*. The development of these infectious clones represents an advancement in the study of USUV, an important emerging pathogen.

## 1. Materials and Methods

### Ethics statement

This study was carried out in strict accordance with the recommendations in the Guide for the Care and Use of Laboratory Animals of the National Institutes of Health. The research protocol was approved by the Institutional Animal Care and Use Committee (IACUC) at Virginia Tech.

### Cell culture

Vero (ATCC CCL-81) and HEK-293T (ATCC CRL-11268) cells were maintained at 37°C with 5% CO_2_ and C6/36 (ATCC CRL-1660) cells were maintained at 28°C with 5% CO_2_. All cell lines were grown in DMEM-5, which consists of Dulbecco’s Modified Eagle’s Medium (DMEM) supplemented with 5% fetal bovine serum (FBS), 1% nonessential amino acids, and 1 mg/ml gentamicin.

### Viruses

USUV strain *Turdus merula* Netherlands 2016 (i.e., “TMN”)(17) was obtained from the European Virus Archives GLOBAL (EVAg). USUV TMN was passaged four times in Vero cells before receipt and one additional time in Vero cells after receipt. USUV strain UG09615 2010 (i.e., “UG”)(18) was a kind gift from the Centers for Disease Control and Prevention (CDC). USUV UG was passaged twice in Vero cells before receipt and once more in Vero cells after receipt.

### Virus sequencing

Next-generation sequencing (NGS) was used to sequence the viral isolates and clone-derived viruses. Libraries were generated by preparing cDNA from total RNA using Maxima H minus reverse transcriptase (RT; ThermoFisher, Waltham, MA, USA) with random nonamer (NNNNNNNNN) primers (Integrated DNA Technologies, Coralville, IA, USA). Second strand synthesis was then performed using Q5 polymerase (New England Biolabs, Ipswich, MA, USA), and libraries were prepared using Nextera XT (Illumina, San Diego, CA, USA). The libraries were sequenced on an Illumina Nextseq 500.

The NGS data were analyzed as previously described with some modifications (19). The pipeline used varied slightly for sequencing the original viral isolates and the clone-derived viruses. For the virus isolates, we first trimmed adapters and low quality reads using BBDuk (20) and aligned the reads to a USUV reference with BWA-MEM (21). We next called variants with LoFreq (22) and filtered the variants present at greater than 50% to a new variant calling file (VCF), which we used to generate a new consensus sequence using bcftools (23). We used the new consensus sequence to re-align the reads and call variants. Fewer reads aligned at the 5’ and 3’ ends of the genomes; accordingly, we performed additional Rapid Amplification of cDNA ends (RACE) sequencing to cover these ends (described below). For clone-derived virus, we used the parental virus consensus sequence as a reference to determine if consensus changes occurred during the cloning process.

To generate the sequences of the 5’ and 3’ ends of the genomes, we used a circular version of RACE based on a previously reported protocol (24). Briefly, viral RNA was extracted using the Quick-RNA Viral Kit (Zymo Research, Irvine, CA, USA), and the RNA was circularized using T4 RNA ligase (New England Biolabs, Ipswich, MA, USA) by incubation at 25°C for two hours followed by a 16°C incubation overnight and enzyme inactivation at 75°C for five minutes. We prepared cDNA from the circular RNA using Maxima RT with a USUV-specific reverse primer (Usu1401r; CAGAGCTGGTAGAACCATGT). We next amplified dsDNA using a forward primer targeted to the NS5 protein (Usu10746f; CAAGCGAACAGACGGTGATG) and a reverse primer targeted to the capsid protein (Usu731r; CGCTTCGAGTGTCTGGTTCT). The amplicons were column-purified and submitted for Sanger sequencing at the Genomics Sequencing Center at Virginia Tech.

### Construction of USUV infectious clones

A Zika virus (ZIKV) infectious clone with a cytomegalovirus (CMV) promoter at the 5’ end and a hepatitis-delta virus (HDV) ribozyme sequence at the 3’ end was used as the backbone to construct the USUV clones. The ZIKV clone was a generous gift from Dr. Andres Merits and has been described previously (25). Viral RNA was extracted using the Quick-RNA Viral Kit (Zymo Research, Irvine, CA, USA) and cDNA was produced using Maxima RT with random nonamer primers. Overlapping fragments were generated using Platinum SuperFi 2X master mix (ThermoFisher, Waltham, MA, USA) and gel purified from a crystal violet agarose gel (26) using the Nucleospin Gel and PCR Clean-up kit (Machery-Nagel, Düren, Germany). The primers used for cloning are presented in Supplemental Table 1. Fragments were assembled using NEBuilder HiFi DNA Assembly Master Mix at a 1:2 (vector:insert) molar ratio. The HiFi assembly was treated with DpnI to remove any residual vector DNA and electroporated into NEBstable electrocompetent cells (New England Biolabs, Ipswich, MA, USA). Cells were plated onto LB agar with 25 µg/mL chloramphenicol and grown at 30°C overnight, and single colonies were used to grow liquid cultures. USUV plasmids were extracted from bacterial cultures using the PureYield Plasmid Miniprep kit (Promega, Madison, WI, USA). The plasmids were screened to determine complete genome assembly using restriction enzyme digestion.

**Table 1.**
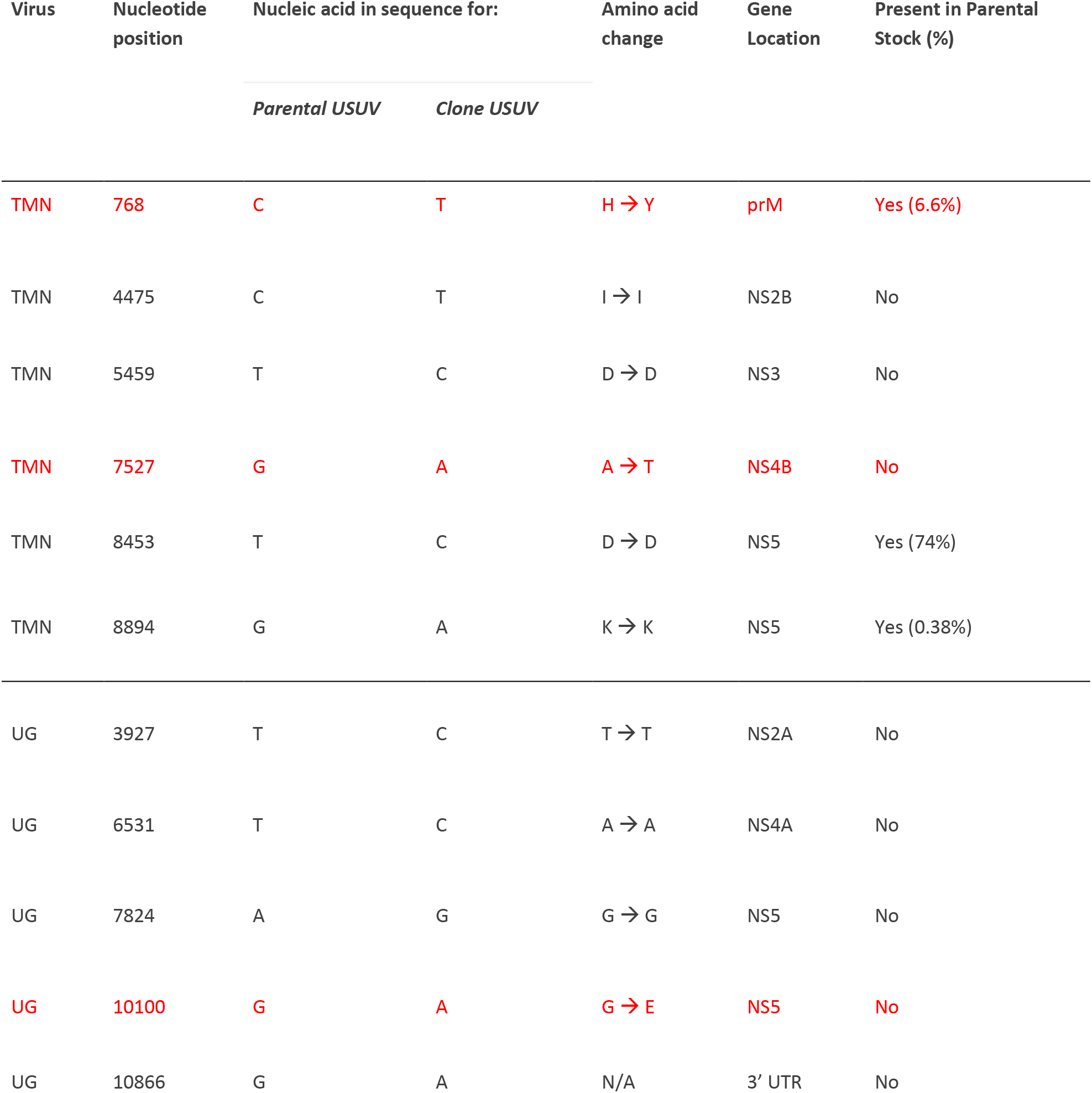
Sequence differences between infectious clone-derived and parental Usutu virus (USUV) strain TM-Netherlands and UG09615. Red text indicates a non-synonymous mutation.

| Virus | Nucleotide position | Nucleic acid in sequence for: |  | Amino acid change | Gene Location | Present in Parental Stock (%) |
| --- | --- | --- | --- | --- | --- | --- |
|  |  | *Parental USUV* | *Clone USUV* |  |  |  |
| TMN | 768 | C | T | H → Y | prM | Yes (6.6%) |
| TMN | 4475 | C | T | I → I | NS2B | No |
| TMN | 5459 | T | C | D → D | NS3 | No |
| TMN | 7527 | G | A | A → T | NS4B | No |
| TMN | 8453 | T | C | D → D | NS5 | Yes (74%) |
| TMN | 8894 | G | A | K → K | NS5 | Yes (0.38%) |
| UG | 3927 | T | C | T → T | NS2A | No |
| UG | 6531 | T | C | A → A | NS4A | No |
| UG | 7824 | A | G | G → G | NS5 | No |
| UG | 10100 | G | A | G → E | NS5 | No |
| UG | 10866 | G | A | N/A | 3' UTR | No |

### Bacteria-Free Cloning

Site-directed mutagenesis (SDM) was performed using a bacteria-free cloning method essentially as previously described (27). Mutated fragments of the USUV TMN and UG clones were amplified by PCR using Platinum SuperFi 2X PCR Master Mix with mutagenic primers (Supp. Table 2) that created 20-25 basepair overlaps. Fragments were assembled using NEBuilder HiFi DNA Assembly Master Mix (New England Biolabs, Ipswich, MA, USA) at a 1:2 (vector:insert) molar ratio. A no assembly control was included that contained the DNA fragments but no assembly master mix to confirm that no residual vector plasmid remained. Assembled DNA fragments were treated with exonuclease I, lambda exonuclease, and DpnI to destroy non-circular DNA and residual plasmid vector, and amplified using the FemtoPhi Rolling Circle Amplification (RCA) kit (Evomic Science, Sunnyvale, CA, USA). The RCA was digested to confirm the correct banding pattern on an agarose gel.

### Rescue of infectious virus

To rescue infectious virus, we transfected 2 µg of the CMV-promoter-containing plasmids or RCAs using JetPrime (Polyplus, Illkirch-Graffenstaden, France) according to the manufacturer’s protocol. Vero cells were used for the original bacteria-based clones and a 70%/30% Vero-HEK-293T cells mix was used for the corrected bacteria-free clones. Virus-containing supernatant was collected when 50-75% of cells displayed cytopathic effects (CPE), aliquoted into working stocks, and stored at -80°C until titration by plaque assay. For bacteria-free cloning samples, only samples with no plaques in the “no assembly control” were used for further studies.

### Growth curves

Vero and C6/36 cells were used to assess viral growth kinetics in 24-well plates. Vero cells were seeded at 8.0 ⨯ 10^4^ cells/mL and C6/36 cells were seeded at 1.0 ⨯10^5^ cells/mL. For infection, a multiplicity of infection (MOI) of 0.01 PFU/cell was used. Virus stocks were diluted in medium, herein called “viral diluent,” consisting of RPMI-1640 with 25 mM HEPES, 1% BSA, 50 μg/mL gentamicin, and 2.5 ug/mL amphotericin B. Each well was inoculated with 50 μl of serially diluted virus mix and all tests were performed in triplicate. Plates were rocked every 10 minutes for 1 hour, and the wells were washed twice with sterile phosphate-buffered saline (PBS). 560 μl of DMEM-5 was added to each well, and 60 μl were immediately collected to measure residual virus. Every 24 hours, 60 μl of viral supernatant were collected, and 60 μl of fresh DMEM-5 were added to each well. Supernatant was collected until 50% CPE was observed, and samples were stored at -80°C until viral titers were determined by plaque assay. The lower limit of detection for plaque assays was 2.26 log_10_ PFU/mL for Vero and C6/36 cells.

### *In vivo* pathogenesis of MAR1-5A3-injected CD-1 and IFNAR^-/-^mice

Three-week-old male CD-1 mice were obtained from Charles River Laboratories (Wilmington, MA, USA). Mice were administered 1 mg of MAR1-5A3 antibody 24-hours before infection via intraperitoneal injection to induce a transiently immunocompromised state. Mice were infected with either the parental strain, infectious clone-derived strain (USUV TMN or UG), or viral diluent as a control, at 2 ⨯ 10^5^ PFU/mouse in the hind left footpad. We weighed and monitored mice daily for fourteen days following infection and bled via the maxillary vein daily for the first four days post-infection to measure viremia. Blood was collected in Microvette 500 Capillary Blood Collection Tubes (Sarstedt Inc., Newton, NC, USA). Serum was collected from collection tubes by centrifugation at 5,000 x g for 10 minutes and stored at 80°C until testing by plaque assay. Studies with alpha/beta interferon receptor-deficient (IFNAR^-/-^) mice were performed in the same manner, except they did not receive MAR1-5A3 antibody, and blood was collected for five consecutive days post-infection (dpi). The IFNAR^-/-^ mice were obtained from The Jackson Laboratory (Bar Harbor, ME, USA) and bred in-house. During the duration of these experiments, mice that lost 15% of their starting body weight or displayed signs of severe disease were euthanized.

### Phylogenetic Tree Analysis

Phylogenetic analysis was performed using the complete coding sequences of various USUV genomes from GenBank NCBI (list available in Supplemental File 1). Multiple alignments of the USUV complete coding sequence genomes were generated using MUSCLE alignment software in MEGA-X (v. 10.1.8). The maximum likelihood method was used for the construction of the phylogenetic tree of 45 USUV, 1 WNV, 1 SLEV, and 1 JEV full coding sequences in MEGA-X using 1,000 bootstraps.

### Statistics

Statistical comparisons were performed in Prism 8 (GraphPad, San Diego, CA, USA). Growth kinetics, qPCR, viremia, and weight-loss were compared using 2-way ANOVA with Sidak’s multiple comparison test. The Mantel-Cox test was used to compare survival in mouse experiments. All negative plaque assay samples were given a value representative of 0.9 plaques on the 10^−1^ dilution.

## 2. Results and Discussion

### 2.1. Usutu Virus Strain Selection and Phylogenetic Analysis

In Figure 1, we present a phylogenetic analysis of 45 USUV and 3 other JEV serocomplex virus full coding sequences using a maximum likelihood method with 1,000 bootstraps. As previously reported, USUV strains cluster into several distinct lineages and are related to other members of the JEV serocomplex (4). For these studies, we selected two recent strains from the Netherlands (TM-Netherlands; herein called USUV TMN) and Uganda (UG09615; herein called USUV UG) to make these infectious clones because of their availability, genetic similarity, recent isolation, and substantial growth differences observed in preliminary studies *in vivo* (Kuchinsky and Hawks, 2020, under review).

**Figure 1.**
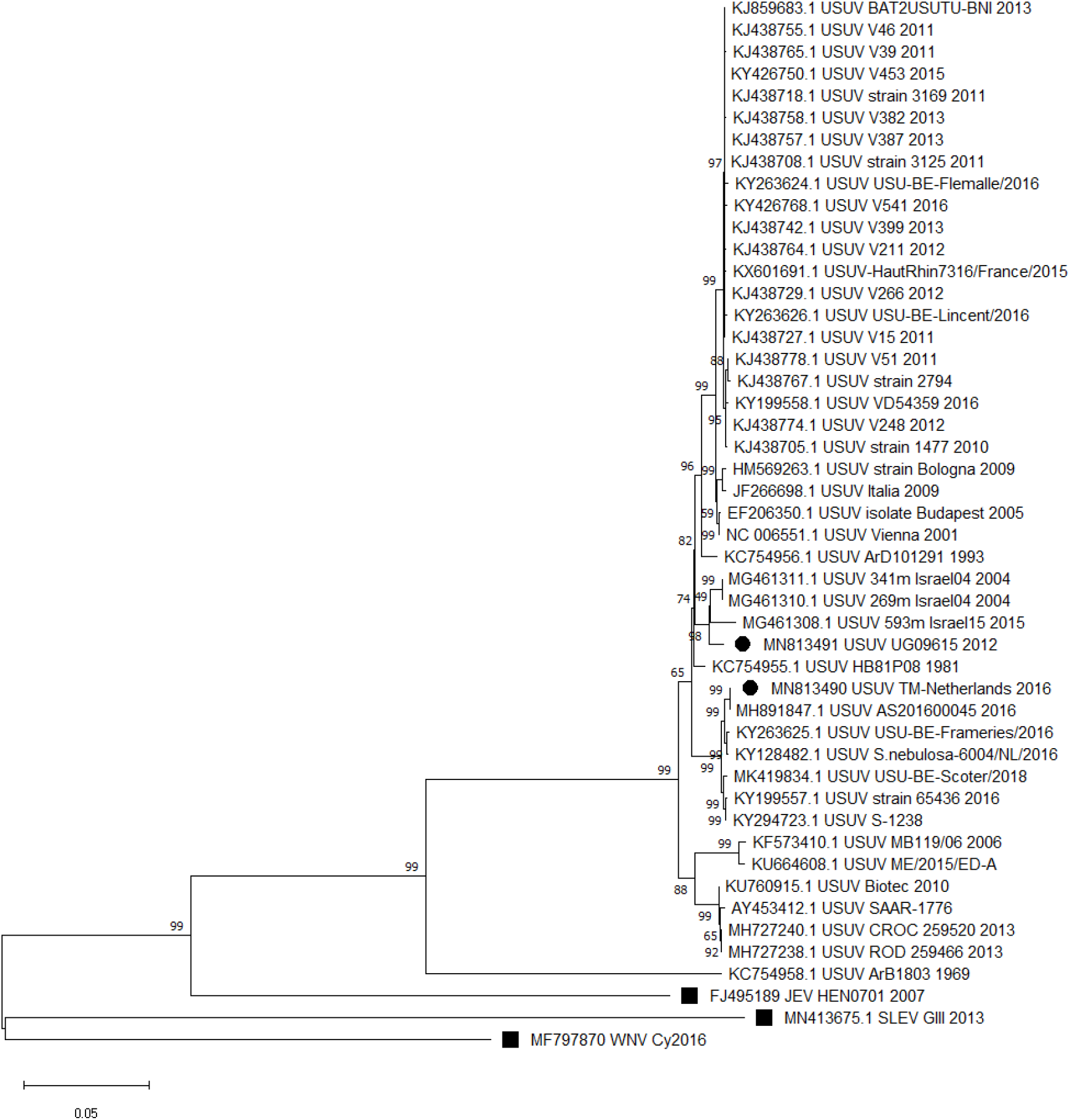
Usutu virus Evolutionary analysis by Maximum Likelihood method. The evolutionary history was inferred by using the Maximum Likelihood method and Tamura-Nei model. The tree with the highest log likelihood (−58247.75) is shown. Initial tree(s) for the heuristic search were obtained automatically by applying Neighbor-Join and BioNJ algorithms to a matrix of pairwise distances estimated using the Tamura-Nei model, and then selecting the topology with superior log likelihood value. The proportion of sites where at least 1 unambiguous base is present in at least 1 sequence for each descendent clade is shown next to each internal node in the tree. This analysis involved 48 nucleotide sequences. There were 10398 positions in the final dataset. Evolutionary analyses were conducted in MEGA-X. USUV strains TM-Netherlands and UG09615 are denoted with black circles. West Nile virus, Japanese encephalitis virus, and St. Louis encephalitis virus are denoted with black squares.

### 2.2. Construction of USUV infectious clones

The infectious clones were derived from parental USUV strains TMN and UG, which were isolated from *Turdus merula* in 2016 and *Culex* mosquitoes in 2010, respectively. These parental strains were sequenced by NGS to create the phylogenetic tree in the previous section (Fig. 1) and for later comparisons to the infectious clones. The backbone for our infectious clones was taken from a pre-existing ZIKV clone containing a CMV promoter and hepatitis-delta virus (HDV) ribozyme sequence. We used a CMV promoter over a bacteriophage promoter since the error rate for RNA polymerase II (RNAPII; 3.9 × 10^−6^ errors per base pair (28)) is roughly 7-13-fold lower than T7 RNAP (2.9-5.0 ⨯ 10^−5^ errors per base pair) (29, 30). Accordingly, CMV-based clones are expected to produce viral stocks with greater homogeneity, limiting the effect of unwanted mutations when studying single-nucleotide polymorphisms and reducing the potential for reversion in vaccines. Finally, despite concerns, splicing of the viral genome does not appear to be a problem in CMV-based clones, even for coronaviruses, which have massive ∼30kb genomes (31). CMV-based infectious clones have been constructed for WNV and many other flaviviruses, validating this approach for the construction of our USUV infectious clones (32, 33).

For clone construction, we used PCR to amplify three overlapping viral fragments and the CMV backbone from a USUV cDNA template and ZIKV infectious clone, respectively. The fragments were then assembled using Gibson assembly and electroporated into NEBstable *E*. *coli* cells.

Infectious virus was rescued by transfecting plasmid DNA into Vero cells, and virus-containing supernatant was collected three days later and titered via plaque assay. The titers for the infectious clone-derived TMN and UG were 1.56⨯ 10^6^ and 1.52×10^7^ PFU/ml, respectively; the titers for parental TMN and UG were 1.58⨯ 10^7^ and 1.50⨯ 10^7^ PFU/ml, respectively.

### 2.3. *In vitro* growth kinetics of parental and infectious clone-derived USUV strains

A clone-derived virus must replicate similarly to the parental virus to be a useful tool. Accordingly, we compared the *in vitro* growth kinetics of the USUV infectious clone-derived viruses and the parental strains in Vero and C6/36 cells following an infection at an MOI of 0.01. The TMN infectious clone-derived strain peaked at an average of 7.62 ± 0.05 log_10_ PFU/mL, while the parental strain peaked at an average of 7.19 ± 0.06 log_10_ PFU/mL on day 2 (Fig. 2A). UG infectious clone-derived (on day 3) and parental (on day 2) viruses peaked at an average of 8.55 ± 0.08 and 8.32 ± 0.10 log_10_ PFU/mL in Vero cells, respectively (Fig. 2B). However, the growth of the TMN infectious clone-derived virus was significantly delayed at 1 dpi (**p≤0.005) and 2-3 dpi (*p≤0.05). Similarly, viral titers of the UG infectious clone-derived virus were also slightly reduced when compared to the UG parental virus at 2 dpi (*p≤0.05). These data indicated that the infectious clone-derived USUV strains were attenuated *in vitro*, likely due to mutations in the viral genome that arose during the construction of the infectious clones.

**Figure 2.**
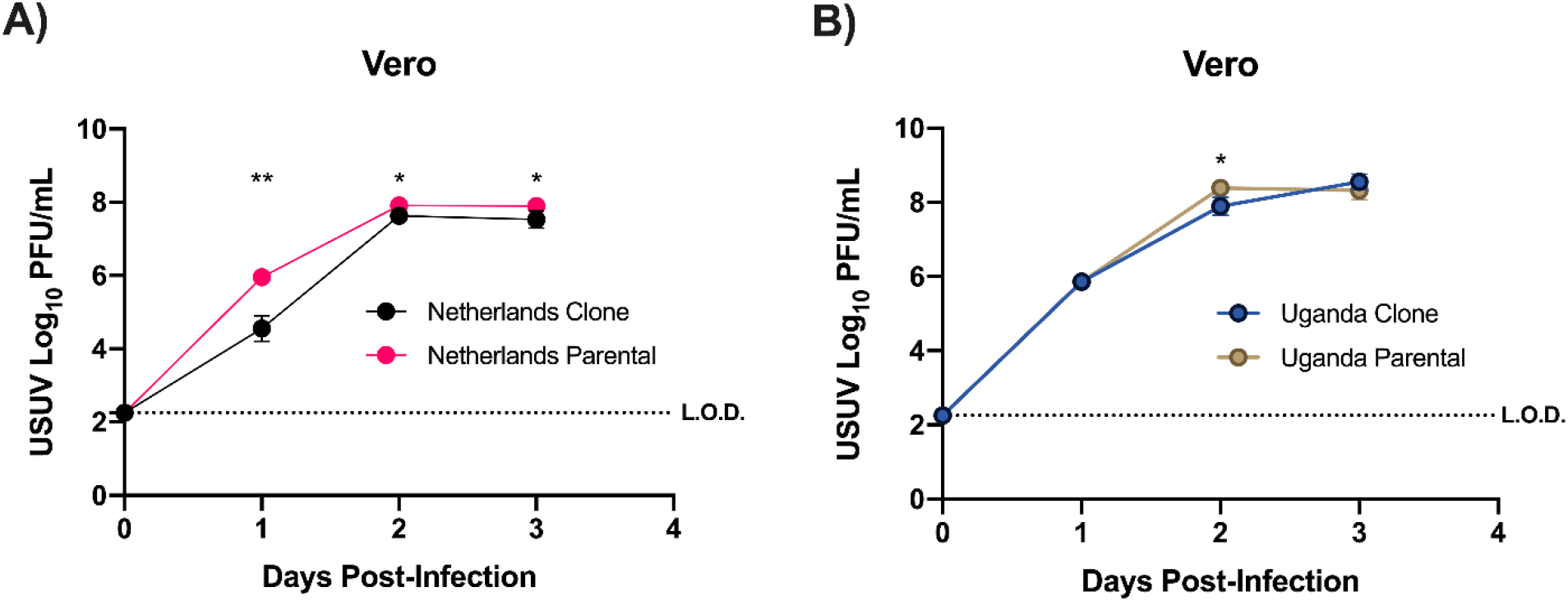
Growth kinetics of USUV TM-Netherlands and UG09615 parental and attenuated infectious clone-derived virus. Growth kinetics were studied in African green monkey kidney cells (Vero) at an MOI of 0.01 PFU/cell. Cells were infected in triplicate with either USUV TMN (A) or UG (B), and supernatant was collected daily starting on Day 0 and tested by plaque assay. Error bars represent standard deviation from the mean. Statistical analysis was done using 2-way ANOVA testing with Sidak’s correction for multiple comparisons. *p≤0.05, **p≤0.01. L.O.D. indicates the limit of detection; the L.O.D. is set at 2.26 log_10_ PFU/mL.

We extracted RNA from rescued virus stocks and sequenced the RNA using NGS to compare the clone-derived virus sequence to the parental virus. We found two amino acid changes present in the USUV TMN infectious clone-derived virus as compared to the parental sequence: a histidine to tyrosine at nucleotide position 768 in the prM gene and an alanine to a threonine at nucleotide position 7527 in the NS4B gene. In the USUV UG clone there was one amino acid change as compared to the parental sequence: a glycine to glutamate at nucleotide position 10100 in the NS5 gene (Table 1). There were also synonymous mutations present in the clones of both strains of USUV. Mutations that were not present in the parental stocks are assumed to have been introduced during PCR, bacterial transformation, or following virus rescue. Some of the mutations were present in the parental stock as minority variants, and the percentage of each is shown in Table 1. The mutation found in the USUV TMN prM region may have altered virion assembly, while the NS4B mutation may have altered IFN antagonism (34). For USUV UG, NS5, the RNA-dependent RNA polymerase is critical for virus replication, and therefore, may have disrupted replication of the genome (34). While interesting, further characterization of these attenuated mutants falls outside the scope of this study. We next sought to correct the non-synonymous mutations towards the goal of generating infectious clone-derived USUV with similar growth kinetics to the parental viruses.

### 2.4. Correction of USUV mutations and *in vitro* growth kinetics of infectious clone-derived viruses

Flavivirus sequences are known to be unstable in bacteria, likely due to cryptic bacterial promoters. This instability can result in attenuating mutations, likely due to the toxicity of flavivirus genomes in *E*. *coli* (35). We hypothesize that the attenuating mutations described above were due to our use of bacteria to construct the clones initially. To negate this issue, we proceeded with the novel method of bacteria-free cloning (BFC) (16), which removes the need for transformation in bacteria by utilizing rolling circle amplification (RCA) to isothermally amplify circular DNA efficiently and rapidly (36). To correct the mutations, we amplified the USUV genome into overlapping pieces using mutagenic primers and the bacteria-derived clones as the template. We assembled these pieces using Gibson assembly to create a closed circular molecule. Notably, we included a control with the DNA fragments but no assembly mix; this was included in all future steps and indicates that no original plasmid template remains following virus rescue. We then amplified the circular template using RCA before rescuing infectious virus by transfection of the RCA product in a Vero/HEK-293T co-culture. Virus was titered by plaque assay and Sanger sequenced to confirm their sequences before further characterization. The passage 0 titers for these infectious clone-derived stocks were 3.80×10^7^ PFU/ml for TMN and 6.20×10^7^ for UG. No plaques were observed on the “no assembly” control samples (data not shown), indicating no contamination from the original clone-derived virus. All mentions of infectious clone-derived viruses from this point on will be the corrected infectious clone-derivatives.

After correcting the mutations present in the infectious clone-derived viruses, we infected Vero and C6/36 cells at an MOI of 0.01 to compare the growth kinetics. No statistical differences were observed between the parental and the infectious clone-derived viruses of USUV TMN and UG in any cell line tested (Fig. 3). All four virus stocks peaked around titers of 8-8.5 log_10_ PFU/mL in Vero cells, consistent with previous results (37). In C6/36 cells, TMN infectious clone-derived virus peaked at an average of 6.13 ± 0.96 log_10_ PFU/mL, and parental TMN peaked at an average of 5.80 ± 0.19 log_10_ PFU/mL. In comparison, UG infectious clone-derived virus peaked at an average of 8.72 ± 0.28 log_10_ PFU/mL, and parental UG peaked at an average of 8.14 ± 0.51 log_10_ PFU/mL. These results indicate that we successfully corrected the attenuating mutations and generated infectious clones with *in vitro* replication kinetics identical to the parental strains. We next aimed to demonstrate the utility of these clones for *in vivo* studies.

**Figure 3.**
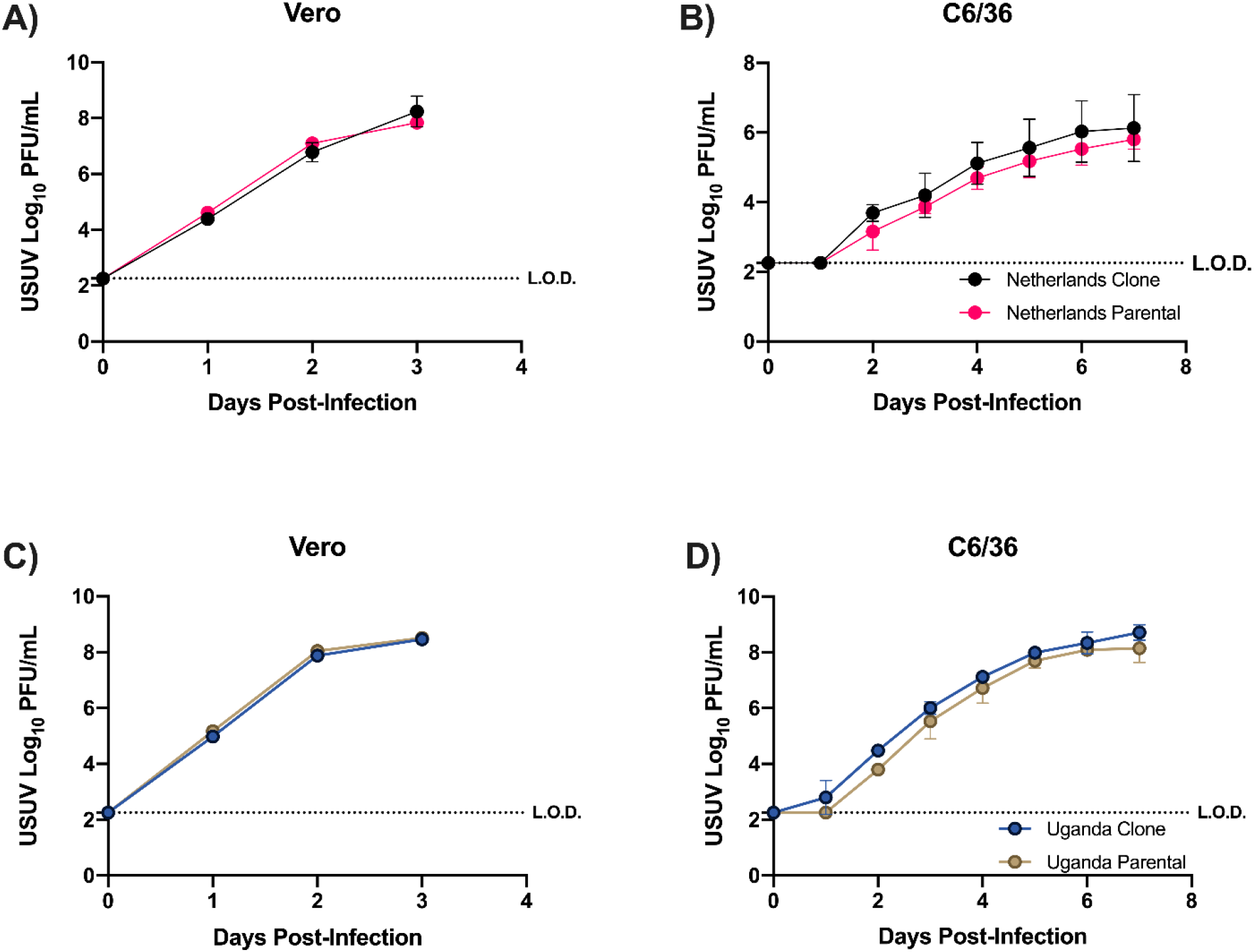
Growth kinetics of parental and infectious clone-derived USUV TMN and UG. Growth kinetics were compared between the corrected infectious clone-derived and parental USUV TMN and UG in African green monkey kidney cells (Vero) (A, C) and *Aedes albopictus* cells (C6/36) (B, D) by infection at an MOI of 0.01 PFU/cell. Cells were infected in triplicate, and supernatant was collected daily starting on Day 0 and tested by plaque assay. Error bars represent the standard deviation from the mean. Statistical analysis was done using 2-way ANOVA testing with Sidak’s correction for multiple comparisons. L.O.D. indicates the limit of detection; the L.O.D. is set at 2.26 log_10_ PFU/mL.

### 2.5. *In vivo* pathogenesis of USUV parental and infectious clone-derived strains

USUV causes sporadic cases of human disease, with severe cases occurring in immunocompromised patients. Therefore, there is an urgent need for a mouse model to study the viral genetic factors contributing to USUV pathogenesis. To that end, we infected transiently immunocompromised CD-1 mice and interferon-alpha/beta receptor-deficient (IFNAR^-/-^) mice with 2×10^5^ PFU of parental or infectious clone-derived USUV (n= 5 mice per group). For CD-1 mice, we administered 1 mg of MAR1-5A3 antibody to transiently block type-1 interferon responses 24 hours before infection. Preliminary studies showed that CD-1 mice infected with USUV without MAR1-5A3 antibody were not susceptible to infection (data not shown); accordingly, it was necessary to use immunocompromised mice for these studies. Our rationale for using MAR1-5A3 was its prior use to render mice susceptible to infection with West Nile virus (38), Zika virus (39), and Ross River virus (40). Furthermore, because the IFN-blockage is transient, mice are expected to be less immunodeficient than genetically deficient mice.

Following infection, viremia in mice infected with TMN parental and TMN infectious clone-derived virus was very similar (Fig. 4A). Infectious virus first appeared on day 2 in some mice, however, viremia was cleared quickly. The TMN clone-derived virus peaked at 2 dpi with an average viral titer of 2.54 ± 0.25 log_10_ PFU/mL. No statistical difference was observed in weight change, and all mice survived (Fig. 4B-C). Due to the low viremia in TMN infected mice, we tested the serum by qPCR as a secondary confirmatory test. Although viremia was low, the qPCR results showed detectable levels of viral RNA from both experimental groups, which peaked at 2 dpi, indicative of viral replication. No statistical differences were observed between the TMN infectious clone-derived strain and the parental strain (Supp. Fig. 1).

**Figure 4.**
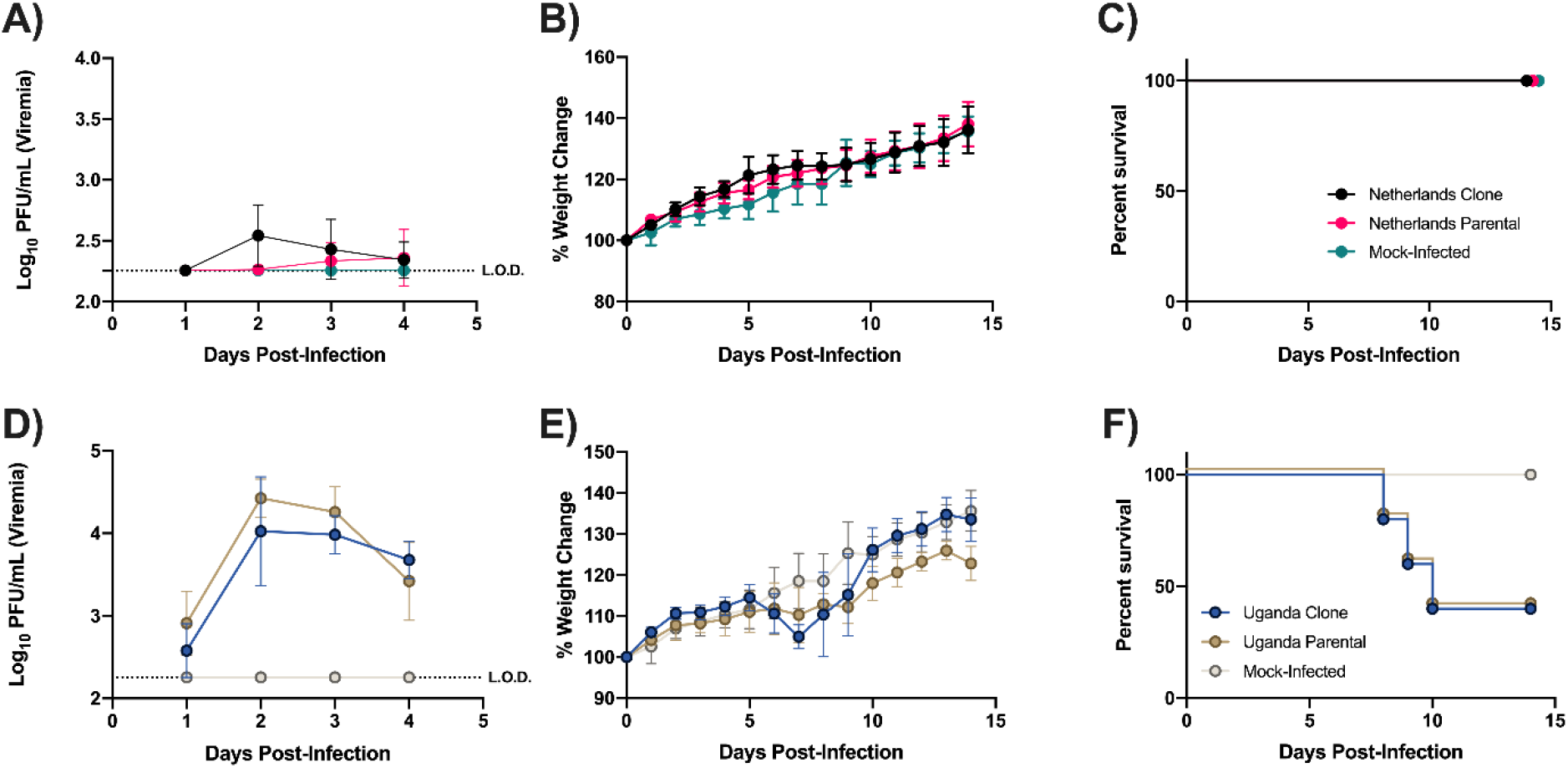
Pathogenesis of infectious clone-derived USUV TMN and UG in MAR1-5A3-injected CD-1 mice. 3-week old male CD-1 (n=5) were administered 1 mg of MAR1-5A3 antibody 24 hours before infection. Mice were infected with 2.0 x 10^5^ PFU of either the parental strain or infectious clone-derived TMN or UG in the hind-left footpad and monitored over time. Blood was collected daily for 4 days post-infection, and the virus titer was determined through Vero plaque assay (A, D). Weights were monitored for 14 days post-infection based on the percent change from starting weight (B, E). Mortality of male CD-1 mice post-infection (C, F). Statistical analysis was done using 2-way ANOVA testing with Tukey’s correction for multiple comparisons. L.O.D. indicates the limit of detection; the L.O.D. is set at 2.26 log_10_ PFU/mL.

Similarly, mice infected with parental UG and infectious clone-derived UG had similar viremia levels (Fig. 4D). In the UG infectious clone-derived group, peak viral titers averaged 4.02 ± 0.65 log_10_ PFU/mL, while the average for the parental group was 4.42 ± 0.06 log_10_ PFU/mL. Weight loss was observed starting 5-7 dpi in both groups, with some mice succumbing to infection at 8-10 dpi (Fig. 4E). There was no statistical difference in weight change over time or in survival between parental UG and infectious clone-derived UG (Fig. 4E-F). We observed a 40% survival rate in both groups (Fig. 4F). Accordingly, in a transiently immunocompromised mouse model, USUV clone-derived and parental viruses behaved identically in replication and pathogenesis.

We next sought to use a more susceptible model — IFNAR^-/-^ mice — which are genetically deficient in type-1 interferon signaling. These mice are highly susceptible to infection by various flaviviruses (39, 41-45). We hypothesized that USUV TMN would replicate more efficiently in these more immunocompromised mice. Following infection, in the infectious clone-derived and parental TMN groups, viremia was detected at 1 dpi and peaked at 4 dpi, at an average of 5.56

± 0.73 and 5.86 ± 0.39 log_10_ PFU/mL, respectively (Fig. 5A). We observed rapid weight loss (Fig. 5B), and all mice succumbed to infection by 6 dpi (Fig. 5C). No differences between the parental and infectious clone-derived TMN viruses were observed for any parameter tested (Fig. 5A-C).

**Figure 5.**
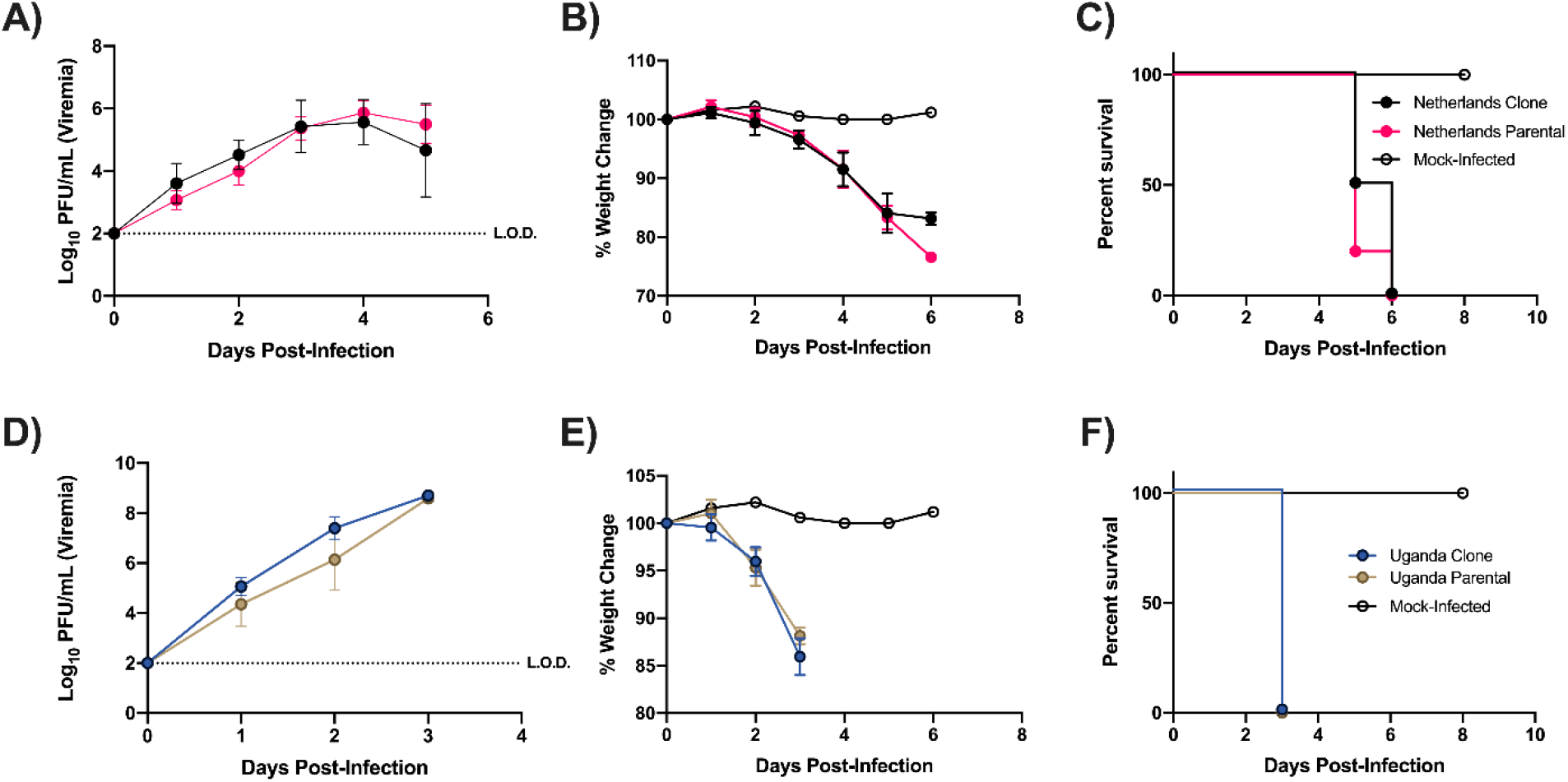
Pathogenesis of infectious clone-derived USUV TMN and UG in IFNAR^(-/-)^ male mice. 3-week old male IFNAR^-/-^ mice (n=5) were infected with 2.0 x 10^5^ PFU of either the parental strain or infectious clone-derived TMN or UG in the hind-left footpad and monitored over time. Blood was collected daily for 5 days post-infection, and virus titer was determined through Vero plaque assay (A, D). Weights were monitored until mortality or euthanasia based on the percent change from starting weight (B, E). Mortality of male IFNAR^-/-^ mice post-infection (C, F). Statistical analysis was done using 2-way ANOVA testing with Tukey’s correction for multiple comparisons. L.O.D. indicates the limit of detection; the L.O.D. is set at 2.0 log_10_ PFU/mL.

Similarly, IFNAR^-/-^ mice infected with parental or infectious clone-derived UG displayed a consistent increase in viremia throughout the experiment. Parental UG had a peak titer average of 8.59 ± 0.73 log_10_ PFU/mL, and infectious clone-derived UG had a peak titer average of 8.70 ± 0.11 log_10_ PFU/mL at 3 dpi (Fig. 5D). We also observed rapid weight loss in the mice, up to 15% of their starting weight (Fig. 5E), and all mice succumbed to infection at 3 dpi (Fig. 5F). Collectively, these data show that these clone-derived viruses behave identically to their parental counterparts.

## Conclusion

In this report, we describe the construction and characterization of USUV infectious clones for two modern strains. We demonstrate that the clone-derived USUV TM-Netherlands and UG09615 replicate similarly to their parental strains and produce similar replication and disease *in vivo* using two mouse models, as well as similar replication in two cell lines *in vitro*.

Interestingly, we serendipitously identified several potentially attenuating mutations that can be further characterized in future studies. These infectious clones may be useful for defining USUV genetic determinants of pathogenesis and may also facilitate the development of recombinant vaccines.

## Acknowledgments

Janet Webster and Kristi DeCourcy.

---

**Supplemental Table 1.**
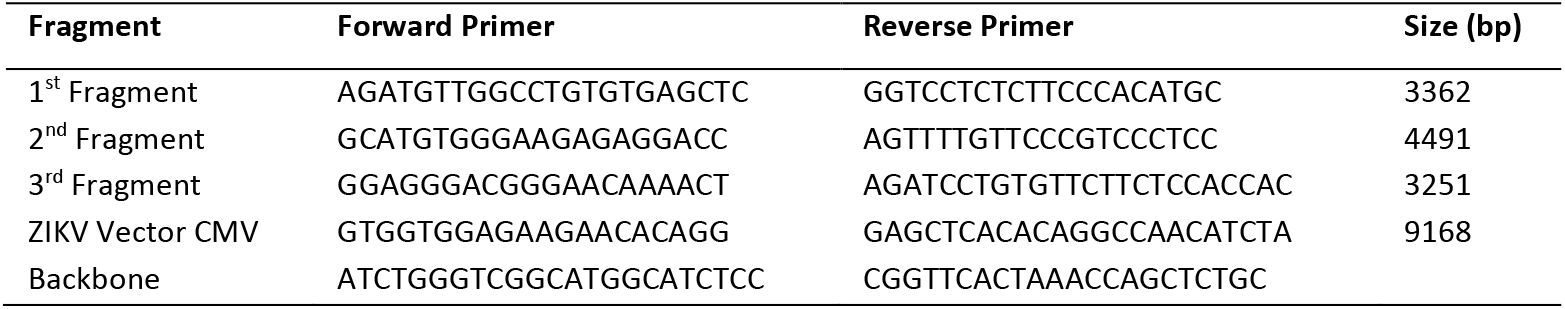
Primer sequences used in the construction of USUV TMN and UG infectious clones.

**Supplemental Table 2.**
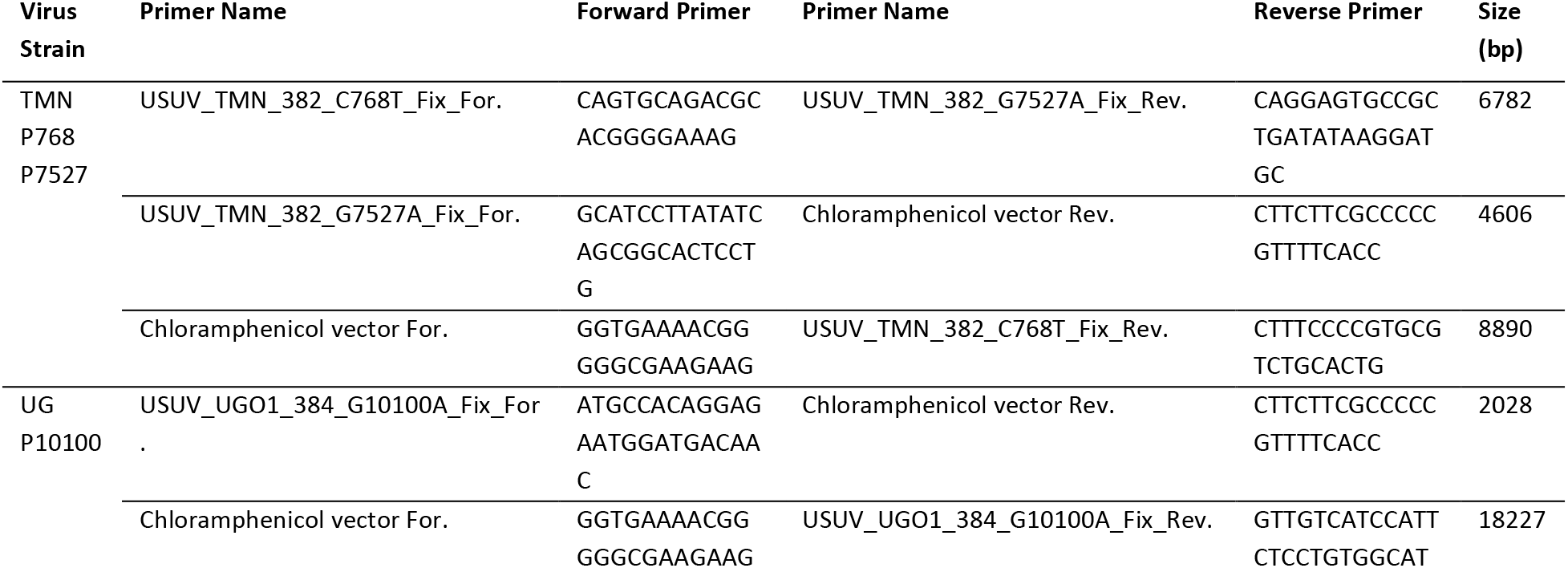
Mutagenic primer sequences used in the correction of attenuated USUV TMN and UG infectious clones.

**Supplemental Figure 1.**
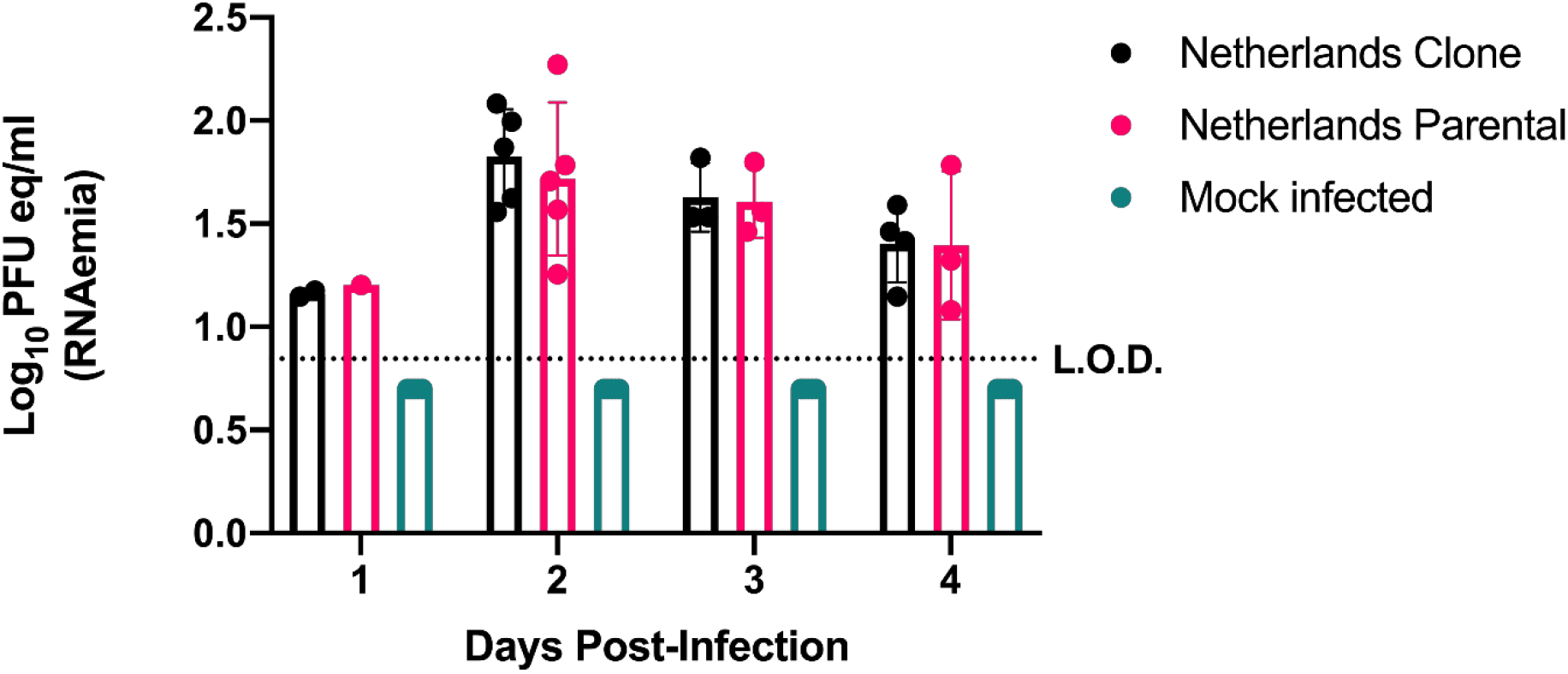
USUV TMN qPCR of serum from infected MAR1-5A3 CD-1 mice. Mice sera were tested via SYBR-green qPCR to determine the presence of viral RNA from TMN-infected MAR1-5A3 CD-1 mice. Serum samples were tested for four days following infection (A). Statistical analysis was done using 2-way ANOVA testing with Sidak’s correction for multiple comparisons. Error bars represent the standard deviation from the mean. RNAemia and the limit of detection (L.O.D.) were calculated using the slope generated from our USUV standard curve. The L.O.D. is set at Cq 35.5 or 0.875 log_10_ PFU eq/mL. Values below the L.O.D. or that did not match the melting temperature of 83.5 were considered negative.

